# Visualisation of gene expression within the context of tissues: an X-ray computed tomography-based multimodal approach

**DOI:** 10.1101/2023.10.09.561491

**Authors:** Kristaps Kairišs, Natalia Sokolova, Lucie Zilova, Christina Schlagheck, Robert Reinhardt, Tilo Baumbach, Tomáš Faragó, Thomas van de Kamp, Joachim Wittbrodt, Venera Weinhardt

## Abstract

The development of an organism is orchestrated by the spatial and temporal expression of genes. Accurate visualisation of gene expression patterns in the context of the surrounding tissues offers a glimpse into the mechanisms that drive morphogenesis. We developed correlative light-sheet fluorescence microscopy and X-ray computed tomography approach to map gene expression patterns to the whole organism’s 3D anatomy at cellular resolution. We show that this multimodal approach is applicable to gene expression visualised by protein-specific antibodies and fluorescence RNA *in situ* hybridisation, offering a detailed understanding of individual phenotypic variations in model organisms. Furthermore, the approach provides a unique possibility to identify tissues together with their 3D cellular and molecular composition in anatomically less-defined *in vitro* models, such as organoids. We anticipate that the visual and quantitative insights into the 3D distribution of gene expression within tissue architecture, by the multimodal approach developed here, will be equally valuable for reference atlases of model organisms development, as well as for comprehensive screens and morphogenesis studies of *in vitro* models.

## Introduction

In the fields of developmental and cell biology, the regional distribution of gene activity can provide insights into the critical processes of an organism’s development and mechanisms causing diseases. Fluorescence microscopy, particularly light-sheet microscopy, is a powerful tool to study gene expression in whole organisms and tissues at high throughput and high spatial resolution (Huisken and Stainier 2009; Krzic et al. 2012; Medeiros et al. 2015; Chatterjee et al. 2018; Keller et al. 2008). Recent developments in the clearing of opaque tissues (Lindsey et al. 2018) and novel geometries in light-sheet microscopy (Daetwyler and Fiolka 2023) enable the visualisation of fluorescent proteins in several centimetres thick specimens. Unfortunately, the specificity to fluorescently labelled tissues also leads to the inability to visualise unlabelled structures. Although general fluorescent dyes such as nuclei and cytoplasm labels or NHS ester (Nanda and Lorsch 2014) can help to portray tissue context, high-resolution imaging of gene expression within a tissue context of a whole organism remains limited to complex and time-consuming multiplexing approaches (Lindsey et al. 2018; Shainer et al. 2023).

The lack of contextual information in fluorescent imaging is typically compensated by imaging of tissue autofluorescence or the use of general dyes. However, this approach takes an extra fluorescence channel calling for multiplexed imaging (Perens et al. 2021; Klevanski et al. 2020) and in the case of autofluorescence, might provide low-level fluorescence. Several multimodal approaches have been suggested to overcome these limitations. Correlative light-sheet and electron microscopy (CLEM) has been used to target and visualise the ultrastructure of bone marrow in zebrafish (Agarwala et al. 2017; Collinson, Carroll, and Hoogenboom 2017). Due to the limited penetration depth of electron microscopy, CLEM is, however, difficult to perform on whole tissues or organisms. Optical coherence tomography has been used to complement fluorescent signals for *in vivo* imaging of a zebrafish embryo (Bassi, Schmid, and Huisken 2015). However, while useful for aligning 4D information or providing general outlines of the sample, optical coherence tomography does not provide sufficient differential contrast of tissues and lacks anatomical detail.

Due to its high penetration depth, X-ray computed tomography is particularly suitable for anatomical imaging of whole organisms with high-throughput (Matula et al. 2021; Weinhardt et al. 2018; dos Santos Rolo et al. 2014; van de Kamp et al. 2018). Based on absorption contrast agents, pan-resolution histology-like, also called histotomography volumes, are obtained in diverse specimens (Ding et al. 2019; Busse et al. 2018; Holme et al. 2014), enabling phenotypic screens (Handschuh and Glösmann 2022; Schoborg et al. 2019) and analysis of pathologies (Kavkova et al. 2021; Eckermann et al. 2020). We therefore decided to combine X-ray tomography and light-sheet microscopy that would achieve simultaneous localisation of fluorescence markers and tissue architecture with pan-resolution in hundreds of micrometre thick specimens.

Here, we show that without modification of typical light-sheet and X-ray microscopes, we can perform a multimodal acquisition that combines fluorescent data with X-ray microtomography, which delivers pan-resolution 3D structural information on whole organisms. We present the cellular precision of this multimodal acquisition and its applicability to targeted protein and mRNA sequences. Furthermore, we demonstrate that our multimodal approach offers possibilities for quantitative analysis of the reorganisation of gene expression upon gene mutation in fish embryos and new insights into the morphogenesis of *in vitro* tissue models.

## Results

### Implementation of a multimodal pipeline

To implement imaging of gene expression within tissue context, we took advantage of a multiview light-sheet microscope (Krzic et al. 2012; Medeiros et al. 2015) and X-ray absorption microtomography optimised for imaging of small model organisms (Metscher 2009; Weinhardt et al. 2018). Both imaging methods allow cellular resolution with short imaging times, enabling high-throughput generation of digital 3D atlases (Matula et al. 2021; Perens et al. 2021; Ding et al. 2019; Pang et al. 2020). As the correlative approach relies on two separate instruments, we optimised sample handling, imaging parameters, and data analysis across a range of fluorescent labelling and gross morphology of samples (Figure 1).

**Figure 1.**
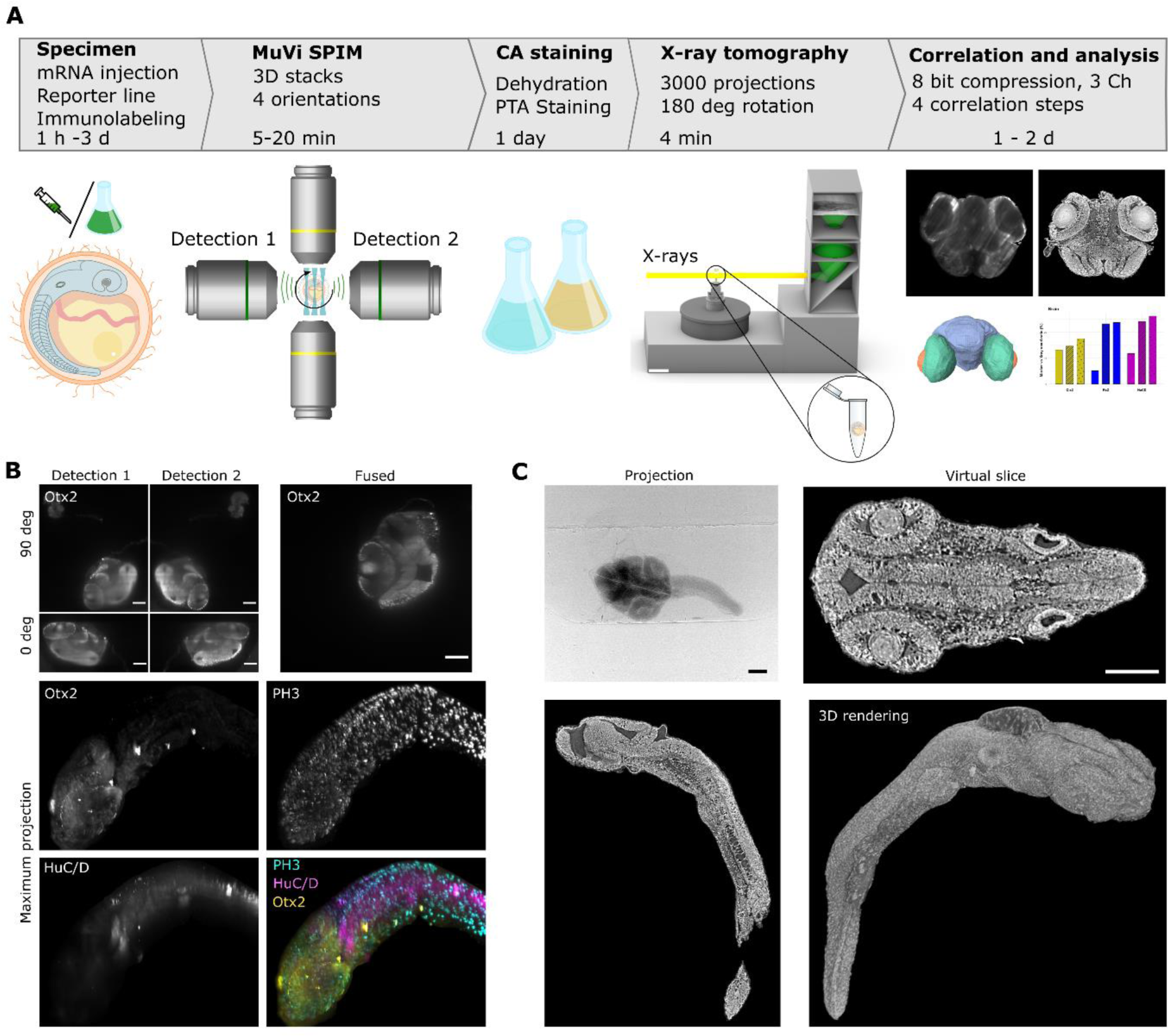
Correlation of light-sheet microscopy and X-ray microtomography into a multimodal pipeline. (A) Scheme representing main stages for the multimodal pipeline. Fluorescently tagged specimens are imaged with the multiview light-sheet microscope (MuVi SPIM), then dehydrated and stained with contrast agent (CA), here phosphotungstic acid for 1 day (depending on the specimen size), imaged with X-ray microtomography, followed by correlation of two modalities and analysis of gene expressions in different tissues. (B) Exemplary fluorescent images were obtained with the multiview light-sheet microscope for detection directions at 0 and 90 degrees rotation of the specimen, and the result of fusion. Maximum projection images from the MuVi SPIM of the medaka embryo (stage 24) immunolabelled with antibodies against Otx2, PH3 and HuC/D proteins. (C) X-ray projection image reconstructed virtual slices and 3D rendering of the same embryo as in panel (B) after staining with phototungstic acid. Scale bars are 100 µm.

Fluorescently labelled specimens were imaged at two orthogonal orientations (0 and 90 degrees) with 2 detection planes at the multiview light sheet microscope (MuVi SPIM) (Luxendo, Light-sheet, Bruker Corporation). Multiview tomographic reconstructions resulted in an isotropic 3D volume with a voxel size of 406 nm and a total volume of about 830 µm × 830 µm × 830 µm (Figure 1B). Due to the high penetration depth of hard X-rays, such volume can be easily imaged with X-ray microtomography; however, it typically suffers from a lack of contrast (Lusic and Grinstaff 2013; Gignac et al. 2016). Therefore, we employed chemical staining with phosphotungstic acid (PTA) to increase the absorption of hard X-rays of specimens as previously reported for different model organisms (Metscher 2009; Weinhardt et al. 2018; Ding et al. 2019; Lesciotto et al. 2020). To implement PTA staining before X-ray imaging, instead of embedding specimens in low melting agarose, we put the specimens on the solid support of 2% agarose, which allows for the exchange of specimens in the MuVi SPIM microscope within several seconds (Supplementary figure 1). For high-throughput imaging, we have imaged PTA-stained specimens with broad energy bandwidth X-ray radiation, resulting in 70 ms exposure time for a single projection, 5 min time for complete specimen acquisition with the 3D volume of 2.5 mm × 2.5 mm × 2.5 mm and isotropic voxel size of 1.22 µm (Figure 1C).

To build up the correlation pipeline, we made use of the image processing package Fiji (Schindelin et al. 2012), mainly the tool for registration and alignment of 3D datasets – Fijiyama (Fernandez and Moisy 2021). In general, all steps can be subdivided into three main parts (Supplementary figure 2): 1) compression of the original data to ease the handling of large datasets; 2) registration of X-ray microtomography to light-sheet datasets; 3) correlation of the original quality data and downstream analysis. The compression of the original data serves two purposes: On the one side, to decrease the volume of the original datasets by cropping (X-ray tomography), rescaling to larger pixel size (light-sheet microscopy) and conversion of 16-bit (light-sheet microscopy) or 32-bit (X-ray tomography) to computationally less demanding 8-bit datasets. On the other side, fluorescent labelling can be sparse and low in its structural content (see Channel 3 in Supplementary figure 2). Thus, combining multiple channels into a single dataset improves registration accuracy. The designed compression steps allowed the downsizing of 72 Gb (light-sheet microscopy, 3 channels, 24 each) and 30 Gb (X-ray tomography) to a few hundred Mb which could be handled without specialised machines for image analysis.

The registration of the datasets was performed at first manually to approximately overlay two datasets and then automatically for accurate registration, typically down to a single nucleus. For manual registration, the use of gross specimen morphology and a few landmarks, such as eye lenses, otoliths, and tail in an embryo (Supplementary figure 3), provided a minimum of 5 datapoints to align light-sheet and X-ray volumes for further automatic registration. After registration, the obtained transformation parameters were applied to the original 32-bit X-ray microtomography data. Notably, performing transformation on an X-ray tomography dataset (and not on light-sheet data) allows to preserve of high spatial resolution of light-sheet data and minimises computational time compared to three channels in light-sheet microscopy. The final correlated multimodal datasets were used to perform segmentation for the digital anatomy of specimens and mapping of gene expression patterns throughout the whole tissue anatomy context (Figure 1A and Supplementary figure 2).

### Cellular precision in the visualisation of gene expression within the context of surrounding tissues

To assess the value of the multimodal imaging, we recorded and superimposed light-sheet and X-ray tomography volumes of stage 24 and 32 medaka embryos (Iwamatsu 2004). Making use of all available filters in the MuVi SPIM microscope, we visualised mitotic cells by anti-phosphorylated histone H3 marker (PH3), newborn neurons using the pan-neuronal marker HuC/D (Good 1995; C.-H. Kim et al. 1996; Park et al. 2000) and forebrain using marker for Orthodenticle homeobox 2 (Otx2) (Acampora et al. 1995; Simeone et al. 1993). The correlation of two imaging modalities showing cellular precision, particularly in the head and the eye regions, is demonstrated by transverse, sagittal, and coronal cross-sections at 1.22 µm^3^ isotropic voxel resolution (see Figure 2A and B, Video 1). The regions at the border of the imaged volume, such as a tail, show artefacts of the registration with the unfaithful position of markers (Figure 2B). Besides, due to dehydration steps applied during the staining procedure for X-ray tomography, the epithelial above the hindbrain was bulging in all datasets (Figure 2). Based on the registration, we found that the dehydration step required for PTA staining results in 20.5±9.5% (n=5) shrinkage of specimens in X-ray tomography compared to light-sheet microscopy. Nevertheless, pan-cellular resolution of X-ray tomography reveals a stereotypic spatial patterning of complex organs such as the developing retina, brain, somites and the developed gut tube along it (Figure 2A, B-B’”). At the outermost layer of the early retina, a single layer of retinal pigment epithelium precursors marked by Otx2-signal in SPIM and can be distinguished by flat cell morphology typical for this cell type in X-ray tomography (Figure 2B” and zoom in Supplementary figure 4). For both stages, that is 24 and 32, Otx2 expression is visible in retinal pigmented epithelium and in progenitors or differentiated photoreceptor cells of the retina (Figure 2, Supplementary figure 5).

**Figure 2.**
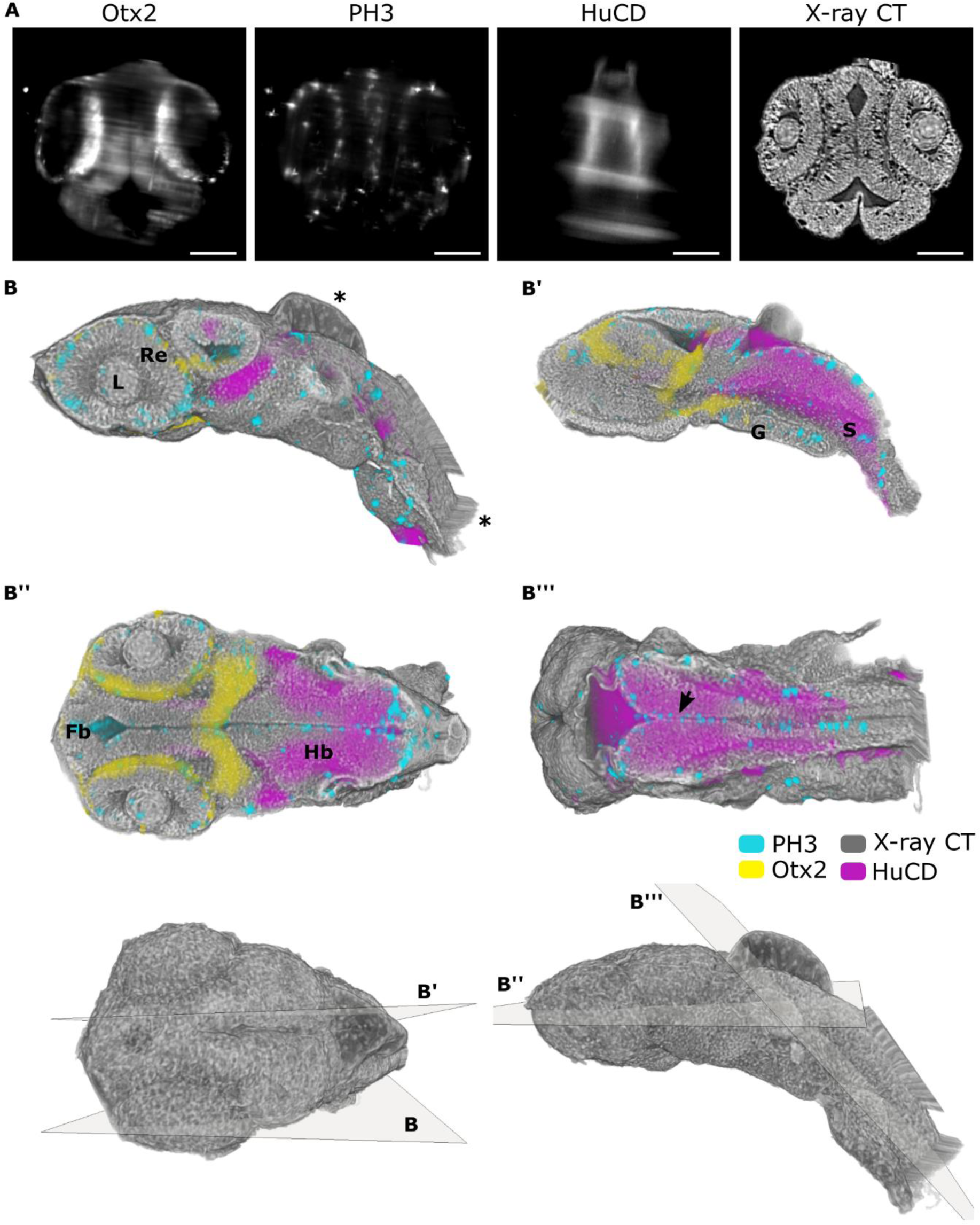
Neuronal genes in the context of tissue anatomy of the medaka embryo visualised by the multimodal approach. (A) Virtual sagittal slices of registered MuVi SPIM and X-ray tomography datasets. From left to right immunolabelling of Otx2, PH3 and HuC/D along with X-ray absorption in stage 24 of medaka embryo. (B) 3D renderings of all contrast modalities visualised with virtual cuts in axial, coronal and sagittal planes. The position and orientation of cuts is shown in the 3D rendering of the whole embryo. Misregistration and bulging of the head region due to dehydration is depicted with *. Organs are labelled as follows: lenses (L), retina (Re), forebrain (Fb), hindbrain (Hb), gut tube (G) and somite (S). Scale bars are 100 µm.

Otx2 expression is also found in the choroid plexus and the most posterior part of the mesencephalon at stage 24 (Figure 2B”, Video 1) with further specification in the hypothalamus and a scattered cell population in the optic tectum by stage 32 (Supplementary figure 5 B””). The newborn neurons labelled by HuC/D expression are located in the hindbrain and the neuronal tube (Figure 2B’”). Several studies have reported that dividing cells in the retina and brain migrate apically towards the periphery of the tissue (Weber et al. 2014; Hevia et al. 2022). Correspondingly, the multimodal imaging shows the localisation of mitotic cells at the outer part of the retina and at the interfaces of the hindbrain and somites which are organ regions unconstrained by any surrounding tissue (Figure 2B, B’” and zoom in Supplementary figure 4).

Due to the difference in contrast between two imaging modalities, X-ray tomography provides information with content beyond the generalised labelling of nuclei or cell cytoplasm. The difference in contrast is demonstrated by the representative sagittal and coronal cross sections of a stage 24 medaka embryo stained with DAPI and N-cadherin (Supplementary figure 6). Notably, X-rays are unique in the ability to visualise whole tissue structures. Furthermore, the difference in X-ray absorption due to varying density of nuclei and extracellular matrix in organs helps to differentiate individual tissues within organs, such as the brain and layers in the retina, enabling quantitative analysis of individual cells and tissues (Ding et al. 2019; Weinhardt et al. 2018).

### Targeted sequence visualisation in combination with X-ray tomography

While visualisation of gene expression via detection of gene product by protein-specific antibody labelling is telling, we further asked whether the same correlative approach could be applied to specimens after fluorescence RNA *in situ* hybridisation (ISH). ISH allows the labelling of selected DNA and RNA strands and, due to such sensitivity and versatility, becomes the most extensively used technique for gene expression analysis (Nath and Johnson 2000). To facilitate the entry of RNA probes and thus increase their hybridisation efficiency, samples undergo several pre-treatment steps. These include methanol treatment to strip membrane lipids and improve permeability (Hoetelmans et al. 2001) and digestion with Proteinase K to remove nuclear, cytoplasmic and extracellular matrix proteins enhancing probe hybridisation (Teng et al. 2017). As RNA ISH is typically performed as an end-point analysis method (Huber, Voith von Voithenberg, and Kaigala 2018), little is known about the preservation of proteins and connective tissue, which are known to be enhanced by PTA staining used in X-ray absorption tomography (Hanly et al. 2023).

After labelling, followed by PTA staining we have observed a variability in X-ray contrast-enhancement of tissue across developmental stages of medaka fish (Supplementary figure 7). For stages 22 and 23 (Supplementary figure 7, panel A), the tissue preservation was comparable to the differential X-ray contrast observed for the X-ray tomography data after antibody staining, see Figure 2. For developmental stages 28 and 33, with the required time of Proteinase K digestion of 20 min, in comparison to 7 min for earlier stages, the differential contrast of tissues is not observed and PTA staining is visible as a superficial layer (Supplementary figure 7, panel B). In this manner, X-ray tomography can potentially be used to analyse tissue preservation after RNA ISH and other sample preparation techniques.

For specimens with preserved tissue morphology after RNA ISH staining, we have used multimodal imaging to map the expression of key transcription factors for early eye and brain development Pax6 and Otx2. We have used simultaneous detection of *Pax6* mRNA (ENSORLG00000009913), using a *Pax6*-specific antisense mRNA probe from the medaka expression pattern database (Alonso-Barba et al. 2016; Henrich et al. 2003; 2005), and antibody staining to detect expression of Otx2 protein on whole-mount medaka embryos as previously described for zebrafish embryos and larvae (He et al. 2020). Consistent with previous reports (Bertrand, Médevielle, and Pituello 2000; Winkler et al. 2000), *Pax6* transcripts were detected in the optic vesicles, in the forebrain at the site of the optic stalk and the neural tube at stage 22 of the medaka embryo (Figure 3). At this stage, the expression of Otx2 is already apparent in the outer layer of the optic cup where the retinal pigmented epithelial is forming. X-ray tomography shows morphologically distinct populations of cells (Figure 3). Accompanying optic vesicle evagination, two morphologically distinct populations of cells are visible in the retina and forebrain. The cells located at the end of the eye field are elongated and form a contiguous layer with the forebrain. The central part of the eye field is filled with cells of distinct morphology and cell density. This morphological transition might result from the neuroepithelial flow observed in the medaka retina teleost during early eye development (Heermann et al. 2015). Such visualisation of cell patterning in relation to gene expression is possible due to the high spatial resolution and histology-like information of this multimodal approach.

**Figure 3.**
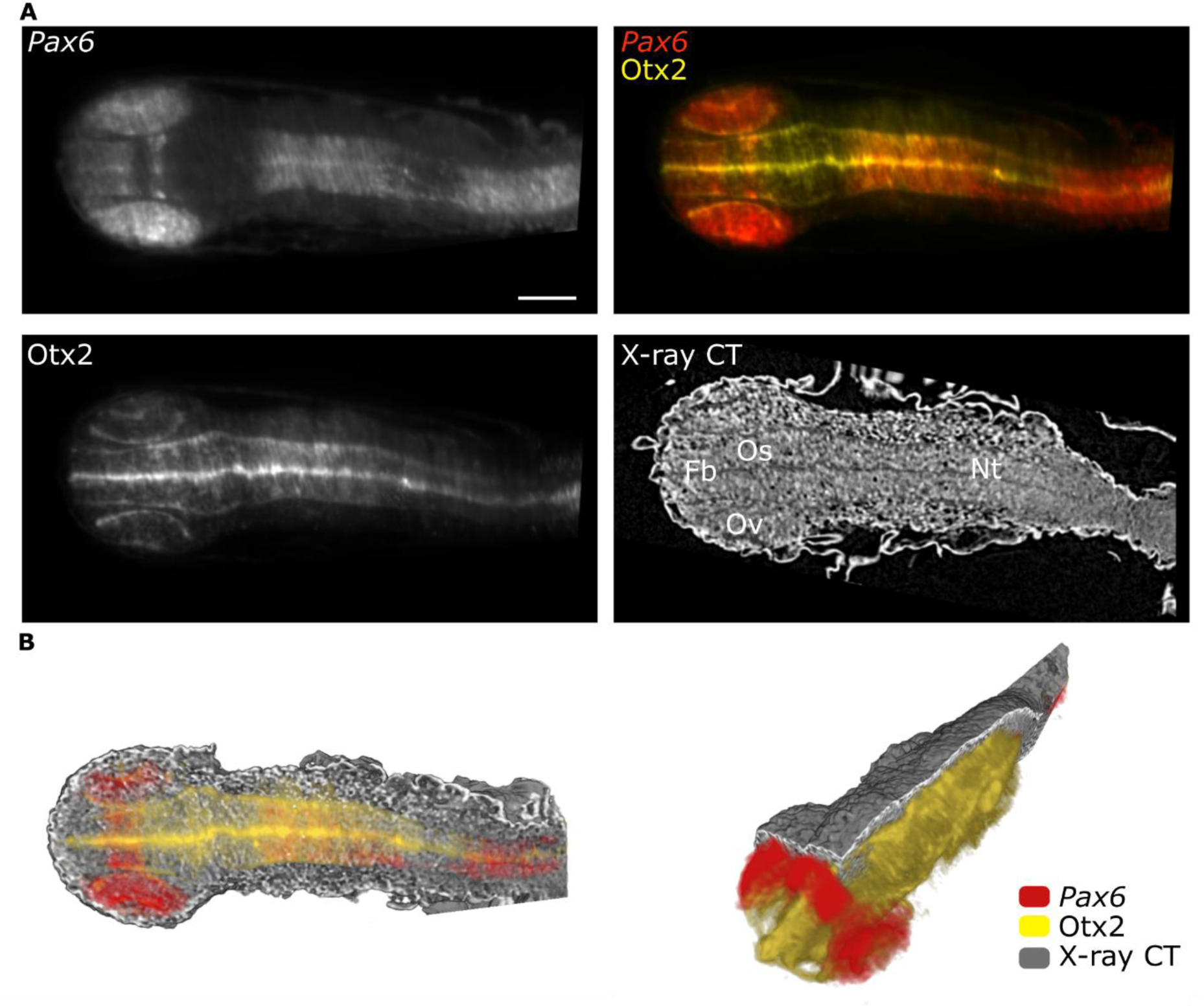
Targeted sequence labelling is correlated with X-ray tomography. (A) MuVi SPIM virtual slices of medaka embryo (stage 22) labelled by fluorescent mRNA *in situ* hybridisation, using *Pax6* mRNA antisense probe and immunolabelled with antibody against Otx2 protein. (B) 3D renderings of all contrast modalities with virtual cuts in transverse and sagittal planes. Organs are labelled as follows: optic vesicles (Ov), forebrain (Fb), optic stalk (Os) and neural tube (Nt). Scale bars are 100 µm.

### Quantitative analysis of phenotypic variations

The multimodal approach can be particularly advantageous when studying morphological mutants. Previously we reported the *Rx3^saGFP^*mutant line, in which retinal progenitor cells in homozygous-mutant embryos fail to migrate laterally to form the optic vesicles (OV), thus mimicking the eyeless mutant (Zilova et al. 2021; Loosli et al. 2003; Rembold et al. 2006). The *Rx3^saGFP^* homozygous mutants express continuous phenotypes, varying from smaller optic cups to the complete absence of eyes (Zilova et al. 2021). Taking advantage of such phenotypic differences and the histological information obtained from X-ray tomography, we asked how the gene expression patterns are altered in various retinal phenotypes. For that, we chose genes marking the main retinal compartments – Rx2 (retinal progenitor cells, ciliary marginal zone, Müller glia and photoreceptor cells), Otx2 (retinal pigmented epithelium, bipolar and photoreceptor cells) and HuC/D (amacrine and ganglion cells), see Figure 4 (Reinhardt et al. 2015; Zilova et al. 2021; C.-H. Kim et al. 1996). Medaka embryos of stage 34 (Iwamatsu 2004) were first subjected to whole-mount immunohistochemistry to detect Rx2, HuC/D and Otx2 proteins and afterwards to X-ray tomography as described above. The resulting volumes were registered onto a combined correlated dataset (Figure 4A). Using the X-ray tomography, we segmented retinas, brain and lenses in the embryo of every group: wild-type sibling (normal eye development), incomplete phenotype (altered optic vesicle evagination or altered optic cup morphogenesis) and complete phenotype (complete absence of the optic vesicle) (Figure 4B). Based on the segmentation, we compared the volumes of each tissue between different phenotypes (Figure 4C-C”). The retina of the incomplete phenotype was much smaller than the retina of a wild-type sibling (Figure 4C). One can also note that the retinal tissue is positioned between two lenses, forming a small optic cup on one side of the embryo (Figure 4B, Incomplete phenotype, retina in green). Such phenotype is caused by the altered migration of retinal progenitor cells from the developing forebrain reported previously (Rembold et al. 2006). The complete phenotype shows an example where such migration was fully arrested, resulting in the absence of optic vesicles (Figure 4B, Complete phenotype). Interestingly, comparing brain volumes (segmented until the otic vesicle in all embryos), we could observe a decrease in the volume for the embryo with incomplete phenotype compared with either wild-type sibling or complete phenotype embryo (Figure 4C’). Moreover, the lens volume decreases along with the increasing severity of the phenotype (Figure 4C”), suggesting the cross-talk between the retina and surface ectoderm which gives rise to the lens in embryonic development.

**Figure 4.**
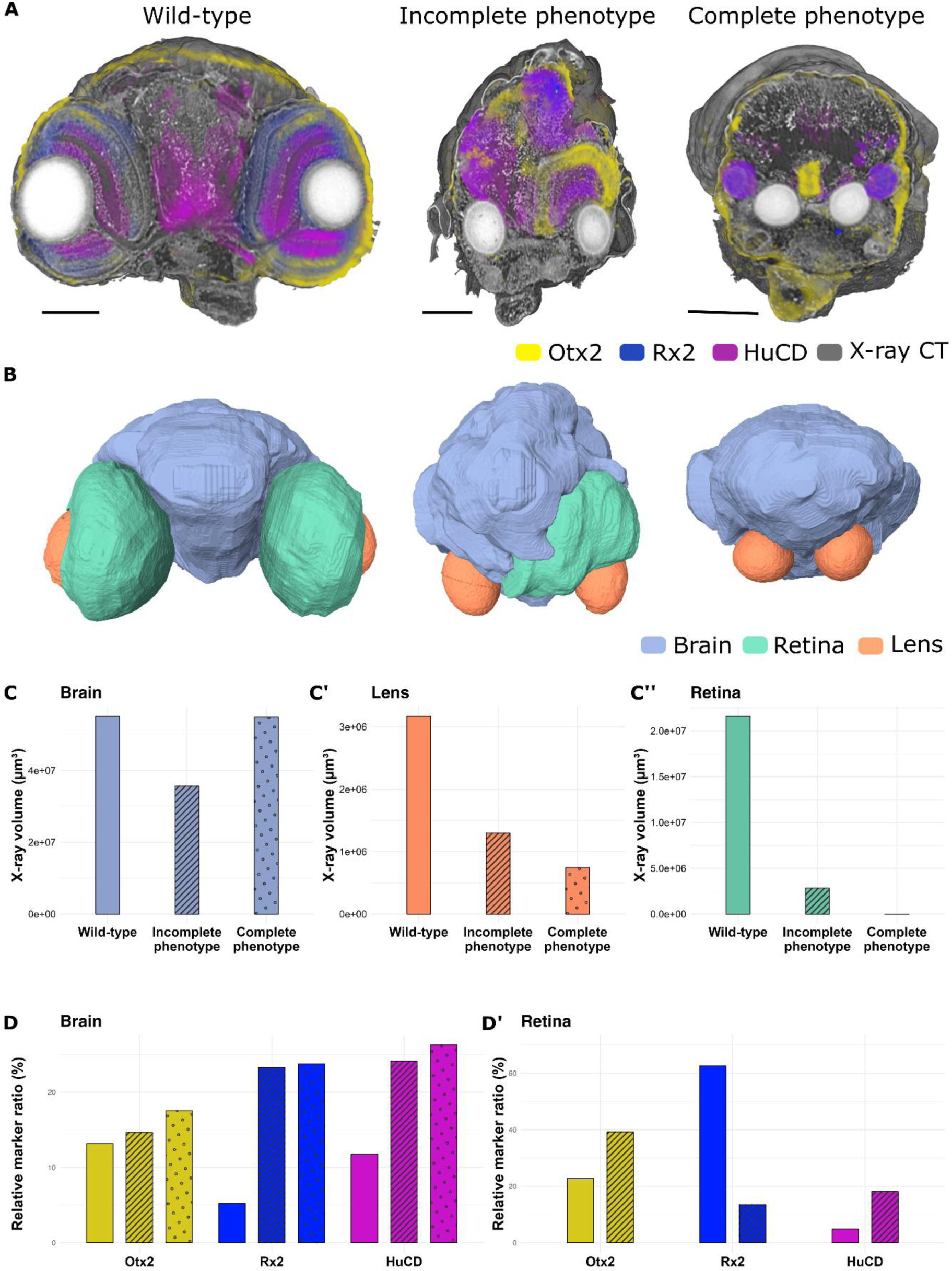
Quantitative analysis of phenotypic variations in *Rx3^saGFP^* mutants. (A) Virtual slices through 3D rendering medaka embryo stage 34 labelled by whole-mount immunohistochemistry Otx2 (yellow), Rx2 (blue), and HuC/D (magenta) markers, imaged by multimodal approach. (B) 3D surface renderings of segmented based on X-ray tomography tissues, that is brain (blue), retina (green), and lens (orange). (C) Brain volume comparison across phenotypes revealed a decrease in the volume in the incomplete phenotype embryo compared with either wild-type sibling or complete phenotype embryo. (C’) Lens volume gets reduced as the phenotype gets more pronounced. (C”) Retinal volumes along the severity of the phenotypes get smaller. (D)-(D’) Quantitative analysis of marker gene expression. In the brain, all the markers rise along the severity of phenotypes. In the retina, Otx2 and HuCD get elevated in the incomplete phenotype while Rx2 gets smaller. Scale bars are 100 µm.

To investigate how the marker gene expression is altered between embryos of various phenotypes, we used automatic Otsu thresholding for all channels within combined labels of retina and brain and obtained the voxel size occupied by either of the markers within the retina or brain. These values were then normalised by the size of the corresponding tissue based on the X-ray tomography (Figure 4, D-D’). Otx2 and HuC/D expression increased in the retina of the embryos with incomplete phenotype when compared to the wild-type sibling, while Rx2 expression drastically decreased. It is likely that proper differentiation of retinal cell types is affected in the incomplete phenotype due to the initially altered architecture of the developing optic cup. Thus, compartments marked uniquely by Rx2 (ciliary marginal zone and Müller glia) may be retinal progenitor cells, present to a lesser extent in the incomplete phenotype retina compared to the wild-type sibling. This would lead to the relative volume increase for the other two markers. In the brain, an increase in the relative ratio was observed for all three analysed markers (Figure 4D’). This can be explained by the fact that, despite a migration arrest of retinal progenitor cells in the *Rx3^saGFP^*homozygous mutants, these cells were still able to obtain the retinal identity while located within the developing forebrain (Zilova et al. 2021).

Such quantitative analysis enabled by the multimodal approach allows for a systematic examination of phenotypic variations at a molecular level in mutants with unknown or continuous phenotypes, like in the case of *Rx3^saGFP^* mutant.

### Multimodal characterisation of 3D organoid systems

The multimodal approach is particularly insightful for organisms that still need to be well described in terms of morphogenesis and 3D cell culture systems, such as explants and organoids. Organoids are valuable for modelling healthy or diseased human tissue and can be highly complex in cell composition (Browne et al. 2017; J. Kim, Koo, and Knoblich 2020). To probe for the complexity of organoids concerning their *in vivo* counterparts, new 3D imaging approaches, which combine the identification of cell types with tissue morphology, are essential.

We have previously reported on the ability of primary pluripotent cells of medaka to aggregate into retinal organoids, recapitulating the medaka-specific pace of development and mechanisms of cell migration during optic vesicle evagination (Zilova et al. 2021). Organoid cultures exhibit significant heterogeneity and variable complexity in cellular composition, calling for new approaches in imaging techniques (Rios and Clevers 2018; Dekkers et al. 2019). We have performed multimodal imaging of fish retinal organoids generated from a transgenic reporter line *Sox2(600bp)::GFP* in which GFP is expressed under the control of a *Sox2* regulatory element to monitor cells with neuronal progenitor identity (Supplementary figure 8). Co-staining of day 2 organoids with the antibodies against GFP, Rx2 and Otx2 proteins (Figure 5) showed that most of the organoid tissue adopted retinal fate with a thick layer of neuroepithelial cells positive for general neuronal marker Sox2 with subdivision to non-retinal and retinal domains based on Rx2 expression (Figure 5 A-B). These non-retinal domains (Rx2^-^) overlap with the expression of the forebrain marker Otx2 suggesting that Rx2^-^ cells adopted anterior neuronal identity as described previously (Zilova et al., 2021). Similar separation of Sox2 and Otx2 domains are observed in the *Sox2(600bp)::GFP* medaka embryos at stage 28 (Supplementary figure 9). In organoids, we have found several domains of Otx2-positive cells with distinct flat morphology. From X-ray tomography, these cells surround the lumen within the retinal organoid and resemble the retinal pigmented epithelial layer seen in medaka embryos (Figure 5C and Supplementary figure 9). Several studies have suggested that lumen plays an active part in tissue morphogenesis (Randriamanantsoa et al. 2022; Tallapragada et al. 2021; Mukenhirn et al. 2023). The complex spatial organisation of the lumen in the fish retinal organoid is clearly visible in the X-ray tomography data (Figure 5D). Though not seen in fluorescence microscopy, the lumen is packed with cellular material, including complex membrane compartments (Figure 5D’). We also found 3 strands integrated into the structure of the organoid (Figure 5B and D). Low X-ray absorption suggests that these are strands of plastics from tips used to pipette cells for aggregation. Visualisation of non-biological material along all cells and extracellular matrix in combination with gene expression patterns might provide an indispensable tool for 3D organoid cultures based on engineered approaches, where cells or cell aggregates are surrounded by biomaterials (Hoang and Ma 2021).

**Figure 5.**
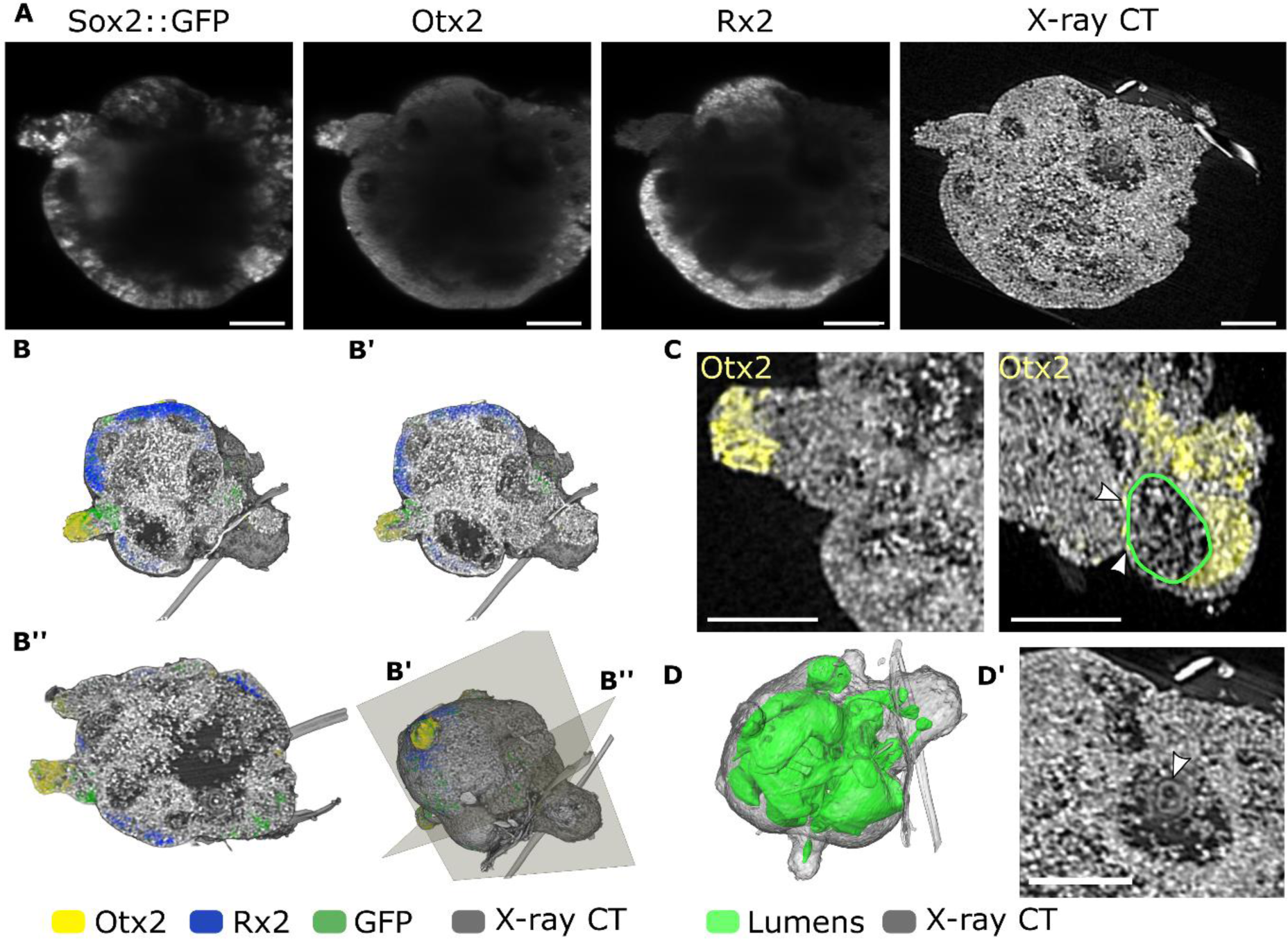
Complex structure of medaka-derived organoids revealed by multimodal imaging. (A) MuVi SPIM virtual slices of day 2 medaka-derived retinal organoids, generated from blastula stage *sox2(600bp)::GFP* transgenic embryos labelled with antibodies against GFP, Otx2 and Rx2 proteins. (B) 3D renderings of all contrast modalities seen in the direction of the volume behind the virtual cuts B’ and B” (the transverse and sagittal planes). (C) Otx2 expression in cells exhibiting different 3D shapes, the lumen is outlined in green. (D) The network of the lumen is shown in green within the organoid. (D’) Cellular content rendered based on X-ray tomography data within lumens is shown in panel (D). Scale bars are 100 µm.

## Discussions

Understanding cellular activity on the molecular level and within the biological context is essential for developmental biology and biomedical research. Previously, to put distinct molecular content back into the context of the surrounding tissue, a combination of bright-field contrast in light-sheet microscopy was exploited by other groups for *in vivo* tomographic reconstructions of overall morphology (Bassi, Schmid, and Huisken 2015). While providing the overall shape of the model organism, higher resolution and tissue sensitivity would significantly empower such correlative imaging approaches. We, therefore combined multiview light sheet fluorescence microscopy with X-ray computed tomography to enable high-resolution 3D imaging of expressed genes in healthy and mutant organisms and in *in vitro* systems.

Compared to other modalities, X-ray tomography offers high spatial resolution and differential contrast of tissues, acting as a hub for histology-like analysis. Through contrast-enhancing sample preparation, X-ray tomography has been used at the synchrotron radiation facilities and with laboratory X-ray tomography scanners to quantitatively assess the morphology of organs, tissues and cell densities within varying model organisms and biological tissues (Ding et al. 2019; Holme et al. 2014; Metscher 2009). Similarly, a high differential contrast of tissues and cell nuclei was achieved in our multimodal imaging of medaka embryos and fish retinal organoids. To avoid shrinkage of tissues due to staining procedure and to simplify the correlation of fluorescence and X-ray signals, light-sheet microscopy could be potentially combined with phase-contrast X-ray imaging (Moosmann et al. 2014; dos Santos Rolo et al. 2014). Low radiation dose of phase contrast X-ray imaging and alternative SPIM geometries are promising approaches to enable *in vivo* multimodal imaging of organisms.

Optical imaging modalities such as light sheet fluorescence microscopy are ideal for targeted visualisation at the molecular level, for example, of expressed genes and DNA/RNA sequences. However, the opacity of biological tissues limits 3D information obtained with fluorescence microscopy to several hundred micrometre-thick specimens. While this information depth was sufficient to analyse the expression of genes in small vertebrate model organism as medaka embryos, the visualisation of larger specimens need further development. To enable imaging of mm thick tissues protocols for clearing of tissues are currently under constant development(Richardson and Lichtman 2015; Lindsey et al. 2018). Clearing protocols based on matching of refractive index, where tissue structure remains close to its native state, will be most promising for multimodal imaging with X-ray tomography.

Our work approaches the ideal of molecular and structural quantitative analysis in 3D for tissue samples and model organisms up to several hundred micrometres in size. The use of X-ray tomography to provide tissue context to the molecular pattern requires no modifications of the existing microscopes, relies on the right order for sample preparation and imaging, and uses existing open-source software to correlate two modalities. Multimodal acquisition provides a complete picture of the composition and molecular identity of tissues, which is especially important when the expression of fluorescence reporters is distributed in different anatomical regions. The ability to track and orient the expression of genes in specific tissues and perform quantitative analysis of their distribution is indispensable for the comparison of multiple samples with abnormal development and morphological defects. The technique is particularly suited to studying emerging model and non-model organisms and *in vitro* 3D tissue models.

## Materials and Methods

### Fish stocks and husbandry

Medaka (*Oryzias latipes*) fish were maintained according to the local animal welfare standards (Tierschutzgesetz §11, Abs. 1, Nr. 1, husbandry permit AZ35-9185.64/BH, line generation permit number 35–9185.81/G-145/15 Wittbrodt). The following lines were used: Cab strain (Loosli et al. 2000) as a wild type and *Rx3^saGFP^* line (Zilova et al. 2021). The *Sox2(600bp)::GFP* reported line was generated for this study as following. The 600nt *Sox2* regulatory element was amplified via PCR from medaka Cab gDNA (forward primer (5’-3’): GAAATAGTAAATATGTACTTAGTATT, reverse primer (5’-3’): CAGGGAAAATTTTAACTTTTCGCTGGG). Plasmid ISceI/MCS-d1GFP-SV40/ISceI was digested with XhoI (NEB) and SmaI (NEB). The *Sox2(600bp)::GFP* plasmid was generated by digesting the primer overhangs of the PCR product with XhoI and EcoRV (NEB) and its consequent ligation into linearised plasmid ISceI/MCS-d1GFP-SV40/ISceI. Injected embryos were raised and maintained at 26°C followed by screening for GFP expression at stages 20-24 on a Nikon SMZ18.

### Fish retinal organoids

The fish retinal organoids were generated as described previously (Zilova et al. 2021). In short, blastula-stage (6 hours post fertilisation) embryos (Iwamatsu, 2004) were collected, dechorionated using hatching enzyme, and washed in ERM (17 mM NaCl, 40 mM KCl, 0.27 mM CaCl_2_, 0.66 mM MgSO_4_, 17 mM HEPES). The cell mass was separated from the yolk and transferred to PBS. After washing twice with PBS, the cell mass was dissociated by gentle pipetting with a 200 μl pipet tip. The cell suspension was pelleted (180 × g for 3 min), re-suspended in retinal differentiation media (GMEM (Glasgow’s Minimal Essential Medium, Gibco Cat#:11710035), 5% KSR (Gibco Cat#:10828028), 0.1 mM non-essential amino acids, sodium pyruvate, 0.1 mM β-mercaptoethanol, 50 U/ml penicillin-streptomycin) and seeded into 96-well plates. The aggregates were incubated overnight at 26°C in an incubator without CO2 control. The following day (day 1) aggregates were washed with retinal differentiation media, transferred to fresh wells, and Matrigel (Corning, Cat#:356238) was added to the media to a final concentration of 2%. From day 1 onward, aggregates were incubated at 26°C and 5% CO2.

### Whole-mount immunohistochemistry

Whole-mount immunohistochemistry was performed as described previously (Inoue and Wittbrodt 2011) with modifications.

Embryos were sorted according to stages 18 to 33 (Iwamatsu 2004) and fixed overnight in 4%PFA in PTW (PBS with 0.05% Tween-20) at 4°C. After fixation, embryos were washed with PTW. Embryos with pigmented retinae were bleached with 3% H2O2, and 0.5% KOH in PTW in the dark. Samples were heated in 50mM Tris-HCl for 15min at 70°C, permeabilised with precooled acetone for 15min at -20°C and incubated in the blocking solution (1% BSA, 1% DMSO, 4% sheep serum) overnight at 4°C. Samples were incubated with primary antibody (1:200) in the blocking buffer for two days at 4°C and washed with PTW 5 times 10 minutes. The following antibodies were used: anti-phospho-histone H3 (rabbit, Millipore, Cat#: 06-570), anti-HuC/D (mouse, Thermo Fisher Scientific, Cat.#: A21271), anti-Otx2 (goat, R&D systems, Cat.#: AF1979), anti-N Cadherin (rabbit, abcam, Cat.#: ab76011). Samples were incubated with secondary antibody (1:750) and DAPI (1:500) or DR (Thermo Fisher Scientific, Cat#: 65-0880-92; 1:1000) nuclear stain in the blocking buffer for 2 days at 4°C and washed with PTW 3 times 10 minutes.

Day 2 organoids were fixed overnight in 4% PFA in PTW at 4°C. Samples were washed in PTW, heated in 50 mM Tris-HCl at 70°C for 15 min, permeabilised for 15 min in acetone at – 20°C, and blocked in 10% BSA in PTW for 2 hr. Samples were incubated with primary antibody (1:500) for 3 days at 4°C. Following antibodies were used: rabbit anti-Rx2 (rabbit, in-house, (Reinhardt et al. 2015), chicken anti-GFP (Thermo Fisher Scientific, Cat#: A10262), goat anti-Otx2 (R&D Systems, Cat#: AF1979). Samples were washed with PTW and incubated secondary antibody (1:750) overnight at 4°C. At the end, samples were washed with PTW and mounted for imaging.

### Fluorescent in-situ hybridisation

Embryos were sorted by stages (Iwamatsu 2004) and fixed in 4% PFA in 2x PTW overnight at 4°C. Fluorescent whole mount *in situ* hybridisations were performed as described previously (Schuhmacher et al. 2011). In brief, samples were rehydrated in the reverse series of MeOH and washed twice for 15 min with PTW. Afterwards, they were digested with 10 µg/ml proteinase K in PTW at room temperature without shaking for the following periods: st.22-23 – 9 min, st.28 – 15 min, st.33 – 75 min. To stop the digestion, samples were rinsed twice in 2 mg/ml glycine in PTW. An additional fixation step was carried out with 4% PFA in PTW for 20 min at room temperature while gently shaken, followed by 5 times for 5 min washing steps in PTW. For the pre-hybridisation, samples were incubated with a hybridisation mix (50% formamide, 5x SSC, 50 µg/ml heparin, 0.1% Tween20, 5 mg/ml torula RNA) at 65°C for 2h. For the probe mix preparation, 10 µl of digoxigenin-labelled *pax6* (ENSORLG00000009913) riboprobe was mixed with 200 µl of the hybridisation mix and denatured at 80°C for 10 min. After pre-hybridisation, the hybridisation mix was replaced by the probe mix. Samples were hybridised at 65°C overnight. Then the probe mix was removed and samples were washed 2x 30 min with 50% formamide in 2x SSCT at 65°C, followed by a washing step in 2x SSCT for 15 min at 65°C, and 2X in 0.2x SSCT for 30 min at 65°C. The following 3 washing steps in TNT (0.1M Tris pH 7.5, 0.15 M NaCl, 0.1% Tween20) were performed at RT for 5 min. Blocking was performed by incubating samples in TNB (TNT with 2% blocking reagent) on a rotator for 2 h at RT. Anti-Dig-POD antibody (1:100) and primary antibody (anti-Otx2, R&D Systems, Cat#: AF1979) in 100 µl of TNB were added to samples. Samples were incubated rotating at 4°C overnight and subsequently were washed 5x 10 min with TNT and rinsed with 100 µl TSA Amplification Diluent. Cy3 Fluorophore Tyramide was diluted 1:50 in TSA Amplification Diluent, and 100 µl of the staining solution was added to the samples. Samples were stained for 2 h at RT without shaking. After the staining, samples were washed 3x 10 min with TNT and incubated with the secondary antibody and DAPI nuclear stain in TNT at 4°C overnight. Afterwards, the samples were washed and imaged with a light-sheet microscope.

### Light-sheet microscopy

3D fluorescent imaging of all specimens was performed on the 16× detection MuVi SPIM Multiview light-sheet microscope (Luxendo Light-sheet, Bruker Corporation). To ensure that the specimens can be used for further imaging with X-rays, instead of mounting specimens with 2% low melting agarose, they were rested on the 2% agarose as shown in Supplementary figure 1. After solidification of the agarose, the tube was further filled with 1x PBS and samples were pipetted inside the tube. The tube was placed vertically such that the specimen fell onto the solid support of agarose.

Two volumes at 0 and 90 rotation each, with 1.6 µm z step size for three channels, that is, GFP 488 nm excitation laser at 80% intensity, 525/50 nm emission filter, 200 ms exposure time, RFP 561 nm excitation laser at 30% intensity, 607/70 nm emission filter, 100 ms exposure time, and far-red 642 nm excitation laser at 50% intensity, 579/41 nm emission filter, 250 ms exposure time, were acquired. The four volumes for each channel were then combined with the Luxendo Light-Sheet microscopy software.

### Synchrotron X-ray microtomography

To improve X-ray absorption of soft tissues, samples were dehydrated in ethanol series of 10%, 30%, 50% and 70% for 1 hour each and stained with 0.3% phosphotungstic acid in 70% ethanol, respectively, for one day at room temperature. After the staining procedure, the samples were washed in 70% ethanol and sealed in polypropylene containers for X-ray tomography.

All specimens were scanned at the IPS UFO 1 tomographic station at the Imaging Cluster of the KIT Light Source. A parallel polychromatic X-ray beam produced by a 1.5 T bending magnet was spectrally filtered by 0.5 mm aluminium to remove low-energy components from the beam. The resulting spectrum had a peak at about 17 keV, and a full-width at half maximum bandwidth of about 10 keV. X-ray projections were detected by a CMOS camera (pco.dimax, 2016 × 2016 pixels, 11 × 11 µm² pixel size) coupled with an optical light microscope (OPtique Peter; total magnification 10×). Photons were converted to the visible light spectrum by an LSO:Tb scintillator of 13 µm. A complete optical system resulted in an effective pixel size of 1.22 µm. For each specimen, a set of 3000 projections with 70 ms exposure time were recorded over a 180° tomographic rotation axis. The 3D volumes were reconstructed using the filtered back projection algorithm implemented in tofu (Faragó et al. 2022).

### Correlation of light-sheet and X-ray microtomography

For the correlation of two modalities in 3D, it is important to find a transformation matrix which would include all degrees of freedom. Because the MuVi SPIM dataset is larger (higher spatial resolution and up to 3 fluorescent channels), the X-ray tomography volume is registered to MuVi SPIM. Before the process of registration, the X-ray tomography volume was compressed from 32-bit to 8-bit depth. Similarly, MuVi SPIM volumes were binned to match the spatial resolution of the X-ray tomography (from 400 nm to 1.22 µm) and converted to 8-bit images. To improve registration of sparse labels all channels of MuVi SPIM were summed up and used for further correlation.

To find the transformation matrix between X-ray tomography and MuVi SPIM, we used the fijiyama plugin designed to correlate datasets from different modalities (Fernandez and Moisy 2021). In short, for rigid transformation, two volumes were rotated and oriented manually, followed by similarities transformation based on points of interest selected in two datatypes as shown in Supplementary figure 2, that is lenses, ears, tail and frontal part of the head. The manual registration was followed by automatic block-matching (number of iterations 20) with similarities, followed by a dense vector field. After the final registration step, all transformation matrixes were merged into one and applied to the original (uncompressed) X-ray tomography data.

### Segmentation, Image analysis and plotting

Segmentation of retinas, brain and lenses was performed in Dragonfly software (Version 2020.2 for Windows, Object Research Systems (ORS) Inc, Montreal, Canada, 2020; software available at http://www.theobjects.com/dragonfly). The fluorescent signal from all SPIM channels was segmented by automatic Otsu threshold using combined retina and brain labels. The resulting mask was then intersected with either the retina or brain label. The values were normalised by the voxel size of the corresponding label. The plots were prepared with RStudio (RStudio Team (2020). RStudio: Integrated Development for R. RStudio, PBC, Boston, MA URL http://www.rstudio.com/) using packages ggplot2 (Wickham 2016) https://cran.r-project.org/web/packages/ggplot2/citation.html, patchwork (Pedersen TL., 2023) https://patchwork.data-imaginist.com/authors.html, ggpattern (FC M, Davis T., 2022) https://coolbutuseless.github.io/package/ggpattern/authors.html, tidyr (Wickham H, Vaughan D, Girlich M 2023) https://tidyr.tidyverse.org/authors.html, grid (Murrell P, 2005) https://stat.ethz.ch/R-manual/R-devel/library/grid/html/grid-package.html. Figures were assembled using Affinity Designer 2 and Inkscape.

## Acknowledgements

This work was supported by grants of the Excellence Cluster “3D Matter Made to Order” (3DMM2O) and Collaborative Research Center (Sonderforschungsbereich) 873 “Maintenance and Differentiation of Stem Cells in Development and Disease” funded through the German Excellence Strategy via Deutsche Forschungsgemeinschaft (DFG), the HIGH-LIFE project (05K2019) via the German Ministry for Research and Education (BMBF), the Carl Zeiss Foundation and the ERC Synergy Grant IndiGene (Number 810172) to JW. V.W. was supported by the German Research Foundation WE 6221/2-1. We acknowledge the KIT Light Source for the provision of instruments at their beamlines and we would like to thank the Institute for Beam Physics and Technology (IBPT) for the operation of the storage ring, the Karlsruhe Research Accelerator (KARA).

## Supplementary information

**Supplementary figure 1.**
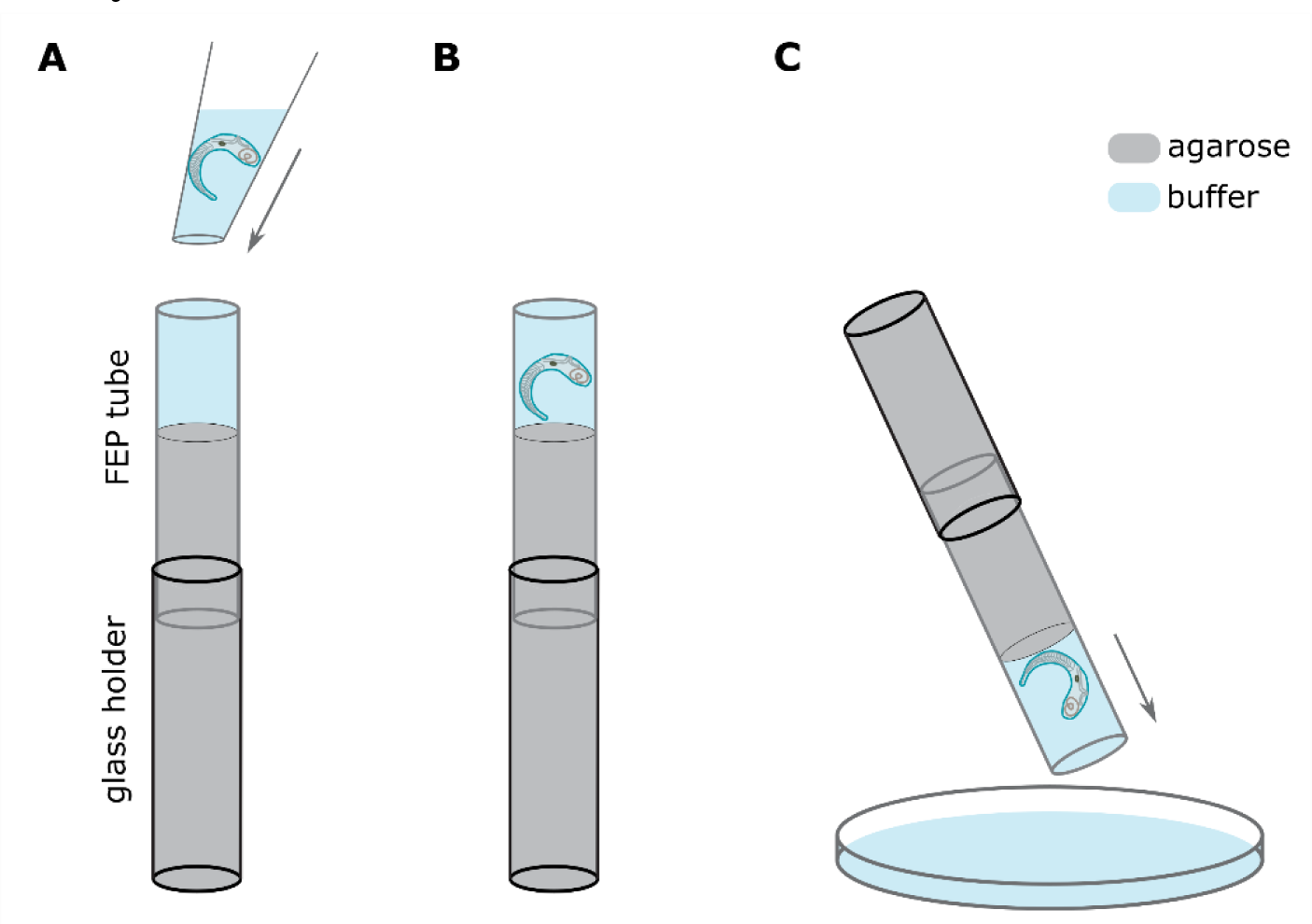
Embedding of specimens for light-sheet microscopy to enable multimodal pipeline. (A) A sample holder for MuVi SPIM consisting of the glass tube and a Fluorinated ethylene propylene (FEP) tube is filled with 2% agarose (in grey) and buffer (in blue). The specimen is pipetted on top with the large opening pipette tip. (B) Due to gravitation, the sample will sink to the bottom of the buffer-filled region, the process can be sped up by gently tapping the tube on the table. (C) The sample is removed by flipping the MuVi SPIM holder into a petri dish filled with buffer. To ensure that the sample is not moving during the acquisition, it is important to select the diameter of the FEP tube matching the gross morphology of the specimen or create a conically shaped agarose mould as previously described (Moosmann et al. 2014).

**Supplementary figure 2.**
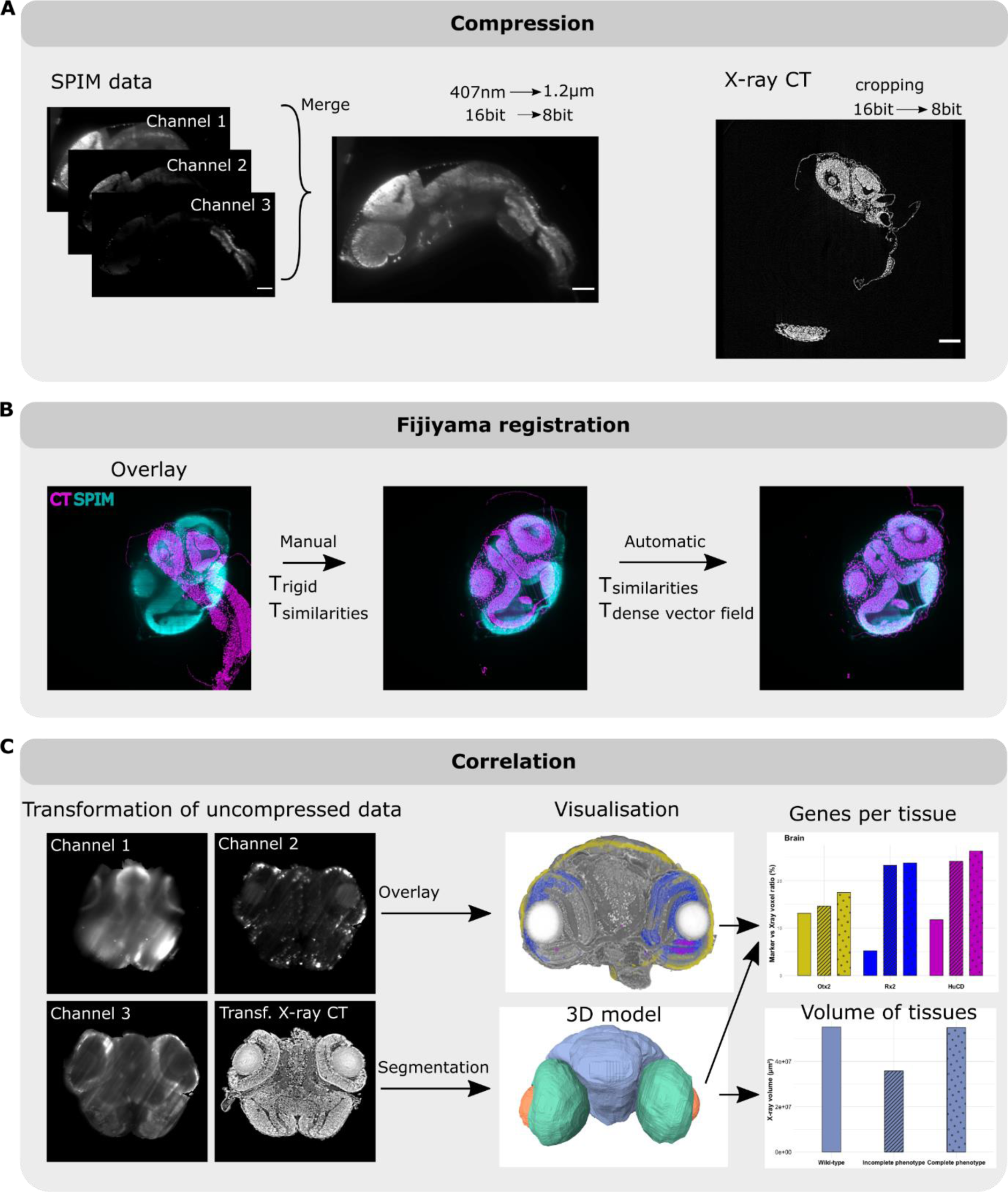
Correlation and analysis of light-sheet microscopy and X-ray microtomography datasets. (A) The first step in a correlation pipeline is compression. All fluorescent signals of MuVi SPIM are merged in one dataset, rescaled to the resolution of X-ray tomography (here 1.2 µm) and compressed to 8-bit. The X-ray tomography is cropped and compressed to 8-bit. (B) The datasets are correlated with the open-source plugin Fijiyama (Fernandez and Moisy 2021). In brief, two datasets are manually oriented to match the overall orientation of the specimen and then rigid transformation is performed on the X-ray tomography dataset. This manual alignment is followed by an automatic block-matching approach with similarities transformation, which would compensate for shrinkage due to dehydration in contrast to deposition for X-ray tomography. Two more automatic registrations are then performed with similarities and dense vector field transformation, resulting in a transformation matrix of X-ray to MuVi SPIM dataset. (C) The transformation matrix is applied to the original X-ray tomography dataset and overlayed with uncompressed individual channels. Based on the tissue structure visible in X-ray tomography, segmentation and quantitative evaluation of tissue morphology and corresponding gene expression are performed. Scale bars are 100 µm.

**Supplementary figure 3.**
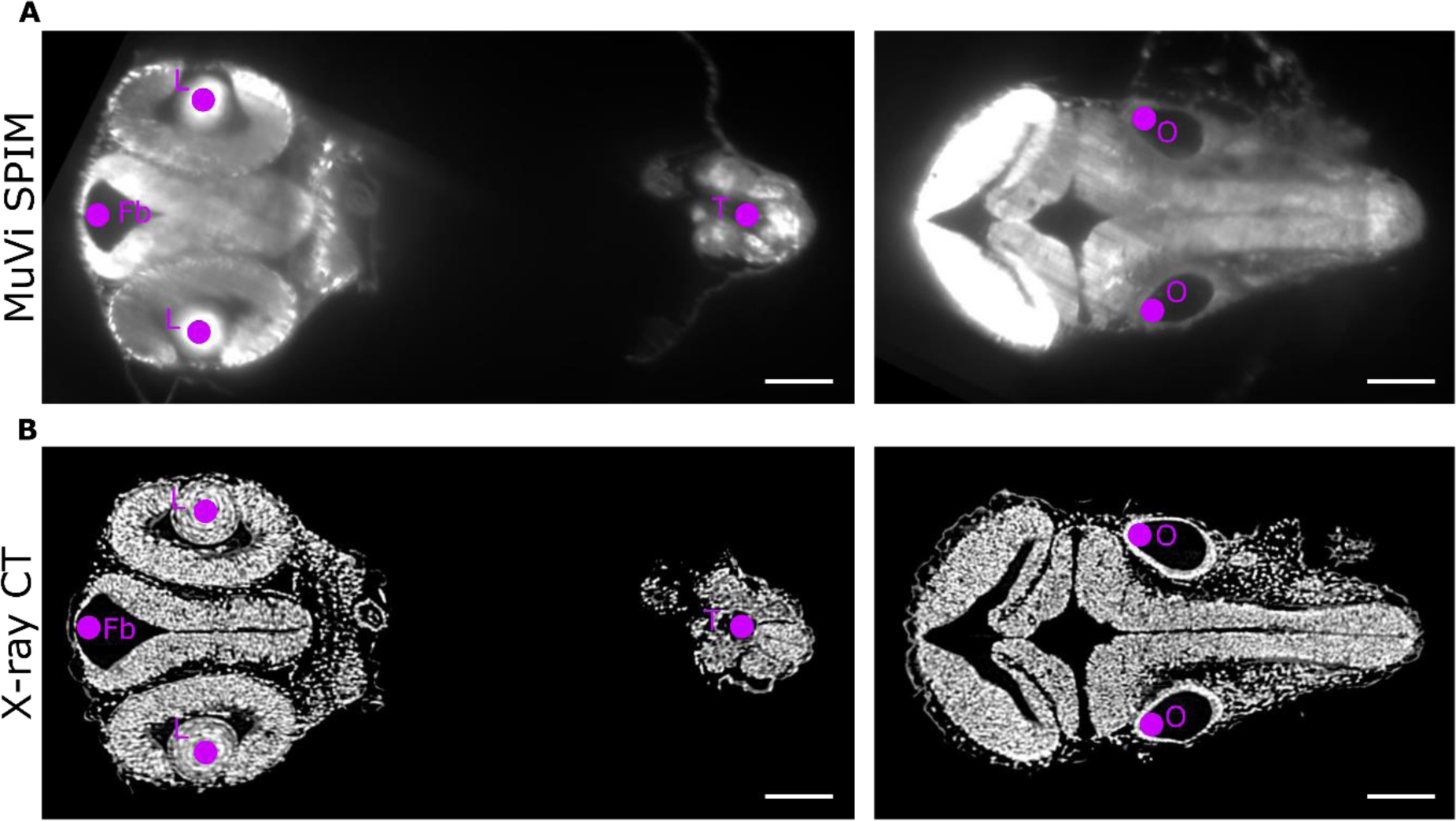
Identification of anatomical features for manual rigid registration. Coronal planes through dataset acquired with the MuVi SPIM (A) and X-ray microtomography (B). A minimum of 5 points are required for rigid registration between datasets. In medaka embryos lenses (L) and otoliths (O) are supplemented by the most anterior part of the brain (Fb) and the most posterior part of the tail (T), specifically the neuronal tube. Scale bars are 100 µm.

**Supplementary figure 4.**
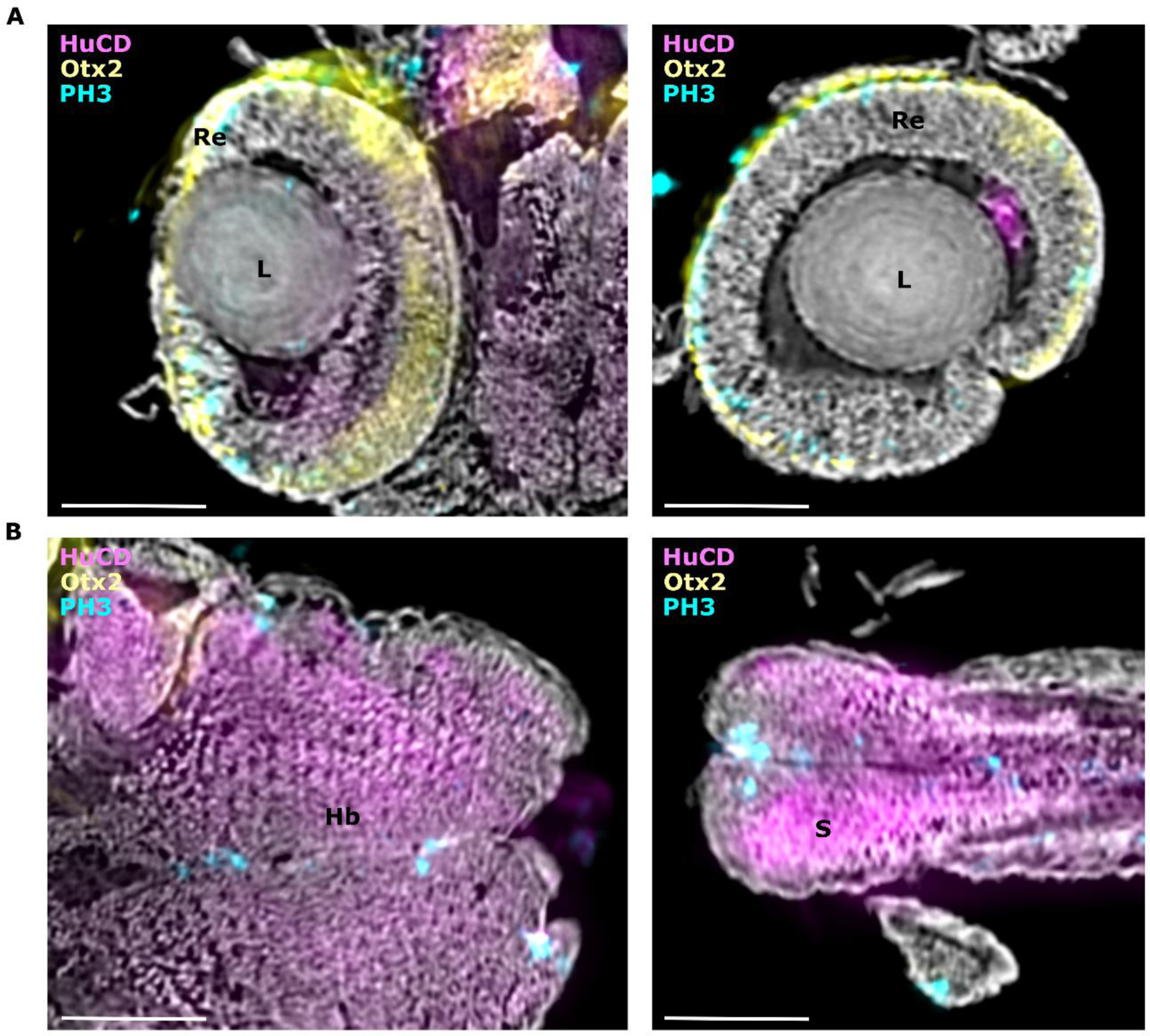
(A) Virtual slices through eye region of cellular precision in multimodal imaging on stage 24 of medaka embryo, shown in Figure 2. (B) Virtual slices through the hindbrain and somite of multimodal imaging on stage 24 of the medaka embryo, are shown in Figure 2. Organs are labelled as follows: lenses (L), retina (Re), hindbrain (Hb), and somite (S). Scale bars are 100 µm.

**Supplementary figure 5.**
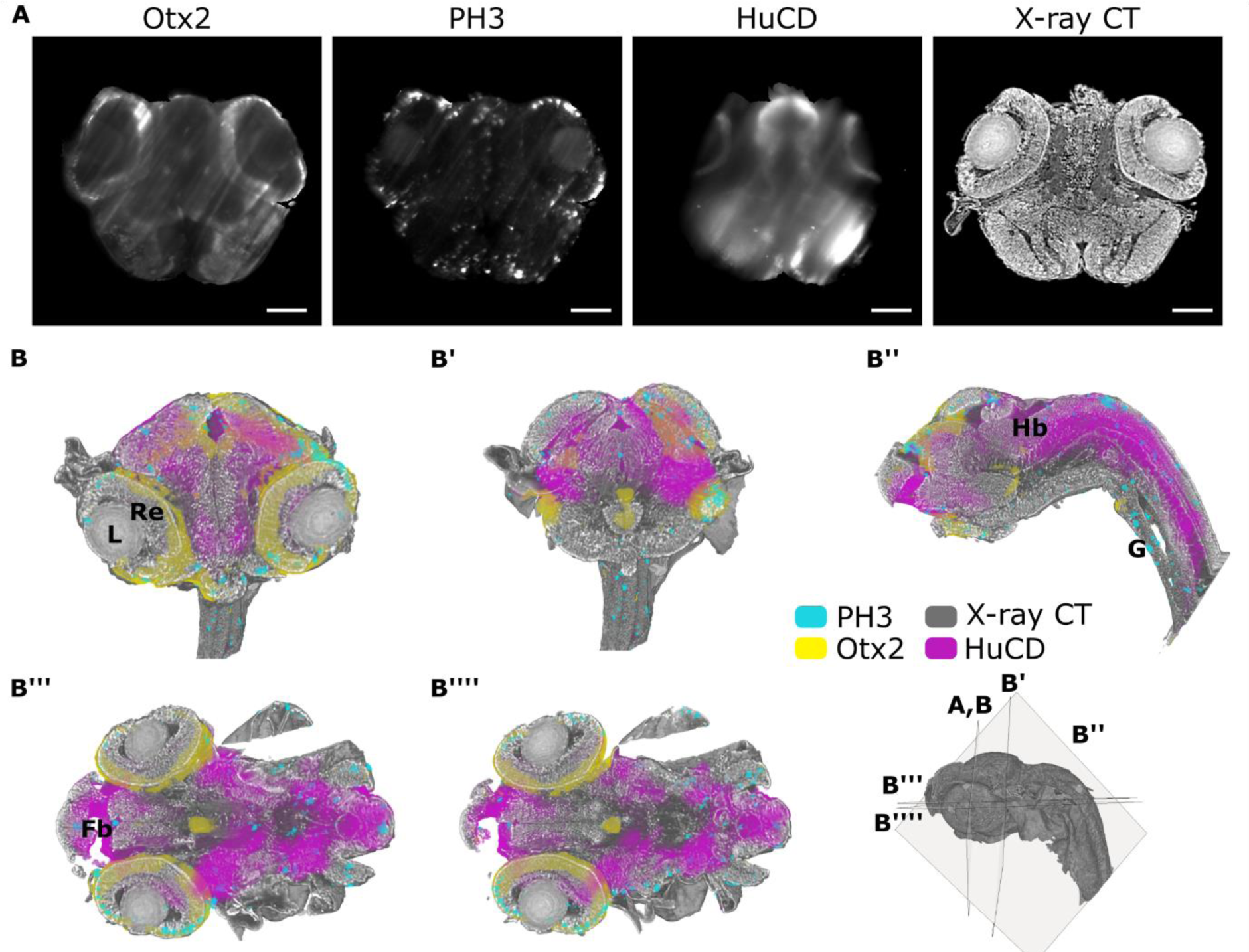
(A) Virtual sagittal slices of registered MuVi SPIM and X-ray tomography datasets. From left to right immunolabelling of Otx2, PH3 and HuC/D along with X-ray absorption in stage 32 of medaka embryo. (B) 3D renderings of all contrast modalities visualised with virtual cuts in axial, coronal and sagittal planes. The position and orientation of cuts is shown in the 3D rendering of the whole embryo. Organs are labelled as follows: lenses (L), retina (Re), forebrain (Fb), hindbrain (Hb), gut tube (G) and somite (S). Scale bars are 100 µm.

**Supplementary figure 6.**
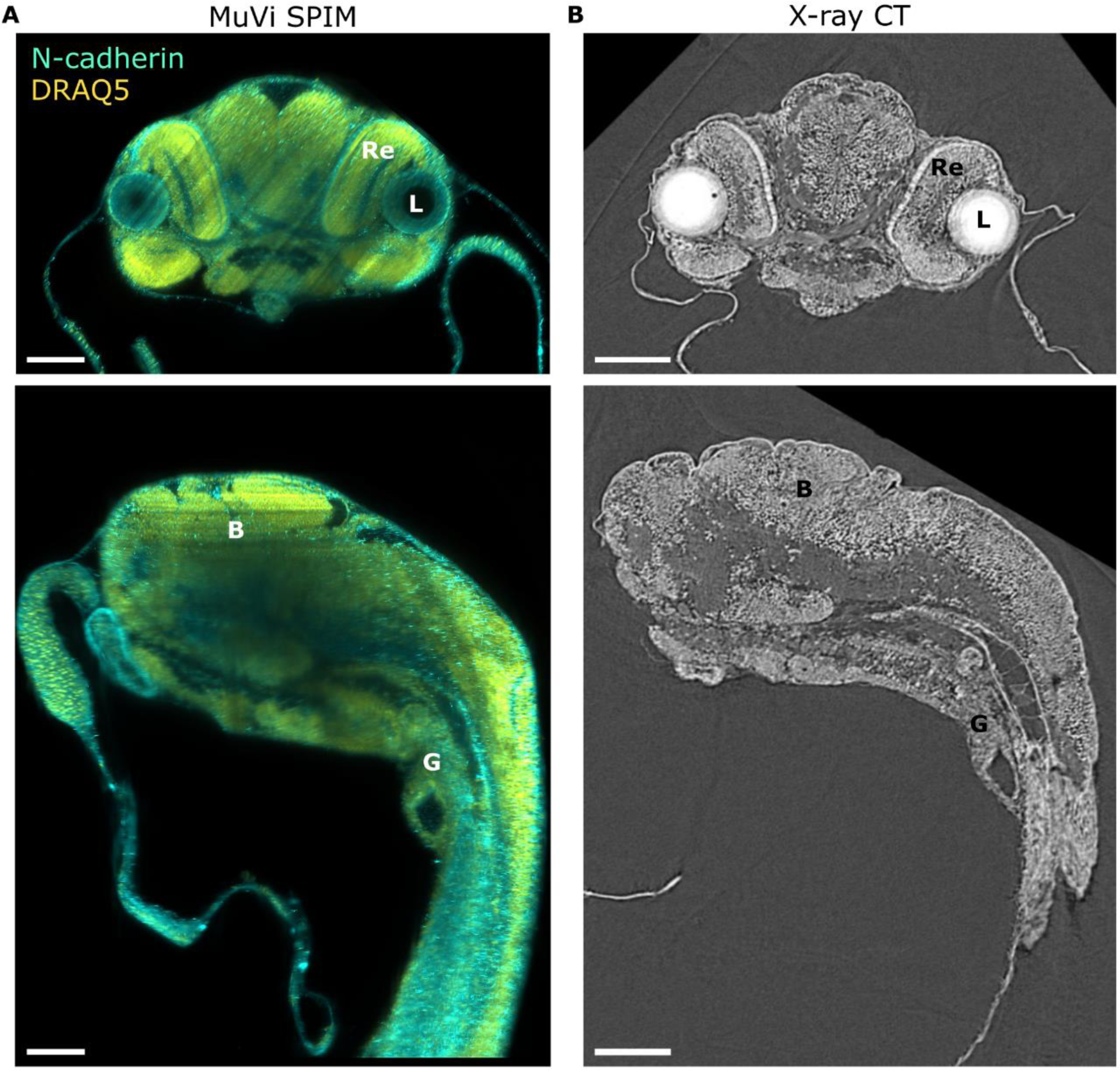
Difference in tissue contrast between fluorescence and X-ray contrast in medaka embryos. (A) Transverse and sagittal virtual slices of medaka embryo (stage 24) stained with antibody against transmembrane protein N-cadherin, co-labelled with nuclear stain DRAQ5 and imaged with the MuVi SPIM. (B) The same as in panel (A) specimen and virtual slices stained with phosphotungstic acid imaged with X-ray tomography. Organs are labelled as follows: lens (L), retina (Re), brain (B) and gut tube (G). Scale bars are 100 µm.

**Supplementary figure 7.**
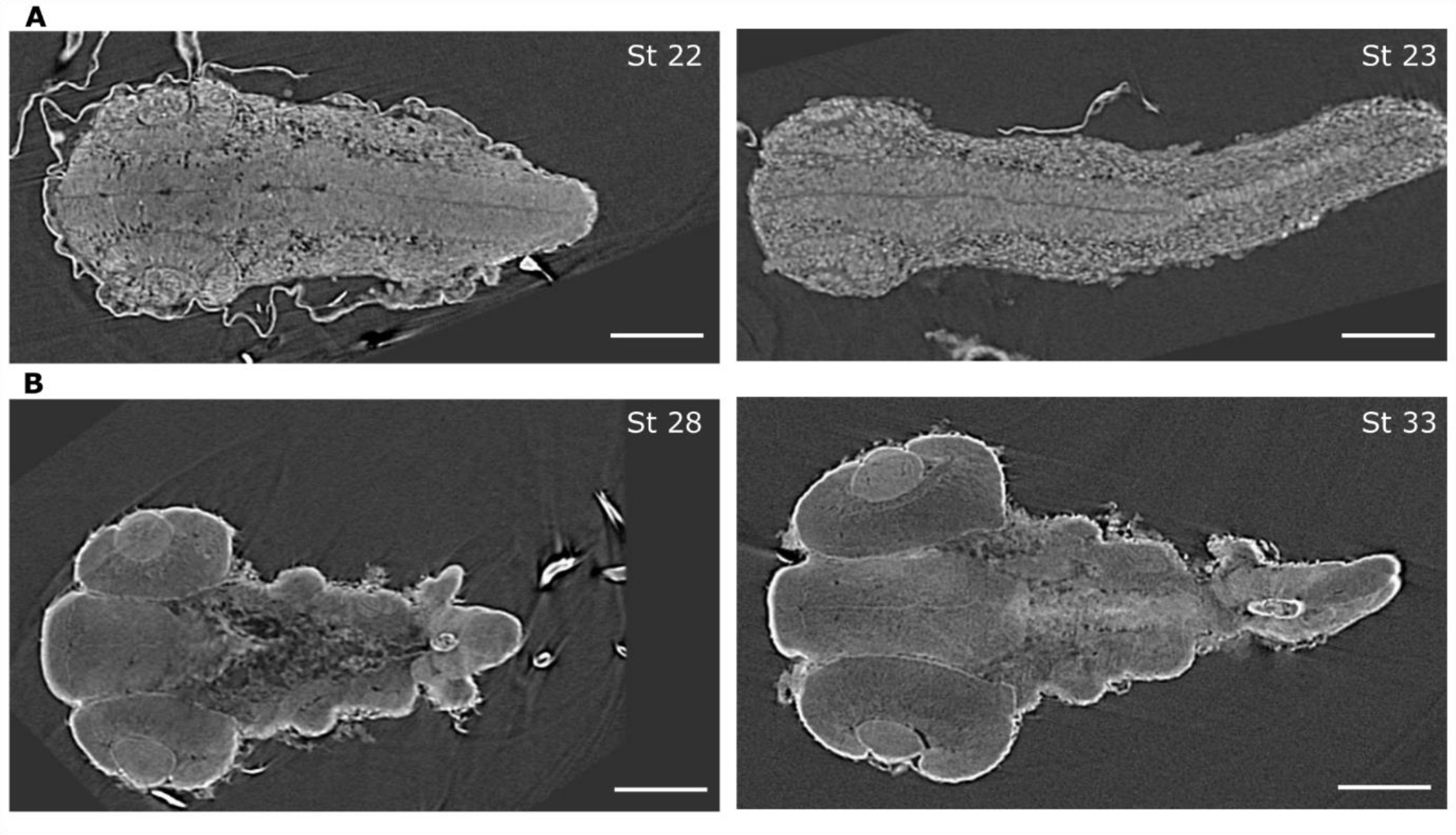
Differences in PTA staining after RNA *in situ* hybridisation for different stages of medaka embryo. (A) Stages 22 and 23 and (B) stages 28 and 33 of medaka embryos with phosphotungstic acid staining after fluorescence *in situ* hybridisation. All specimens were stained with the same reagents and under the same conditions. Scale bars are 100 µm.

**Supplementary figure 8.**
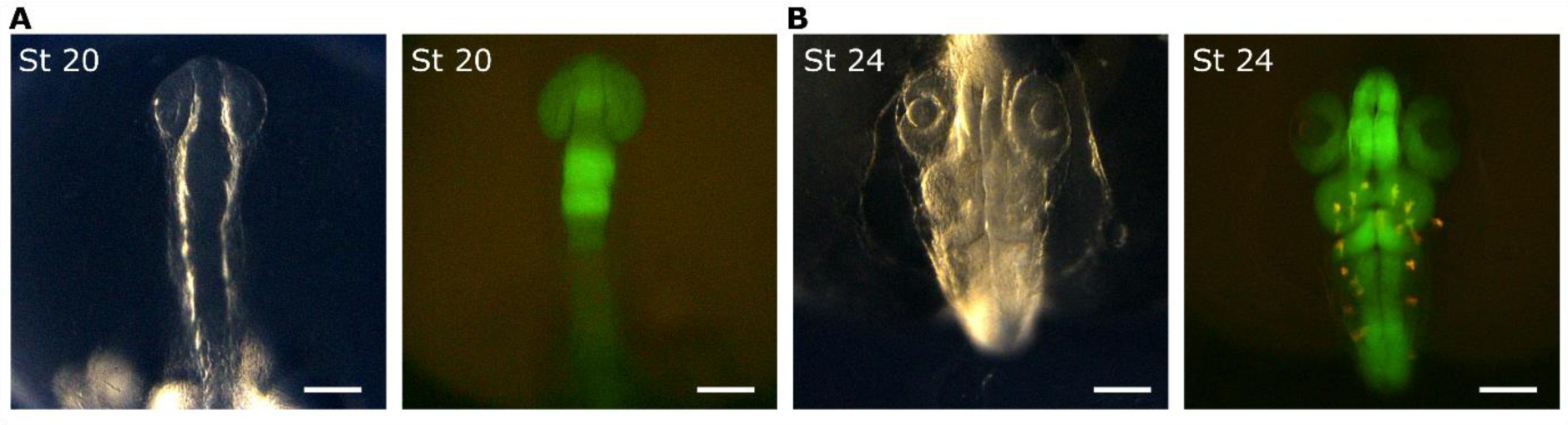
Characterisation of *Sox2(600bp)::GFP line*. Bright-field and fluorescence images of *Sox2(600bp)::GFP line* and corresponding sox2 domain of GFP in the developing embryos at stages 20 and 24. Scale bars are 100 μm.

**Supplementary figure 9.**
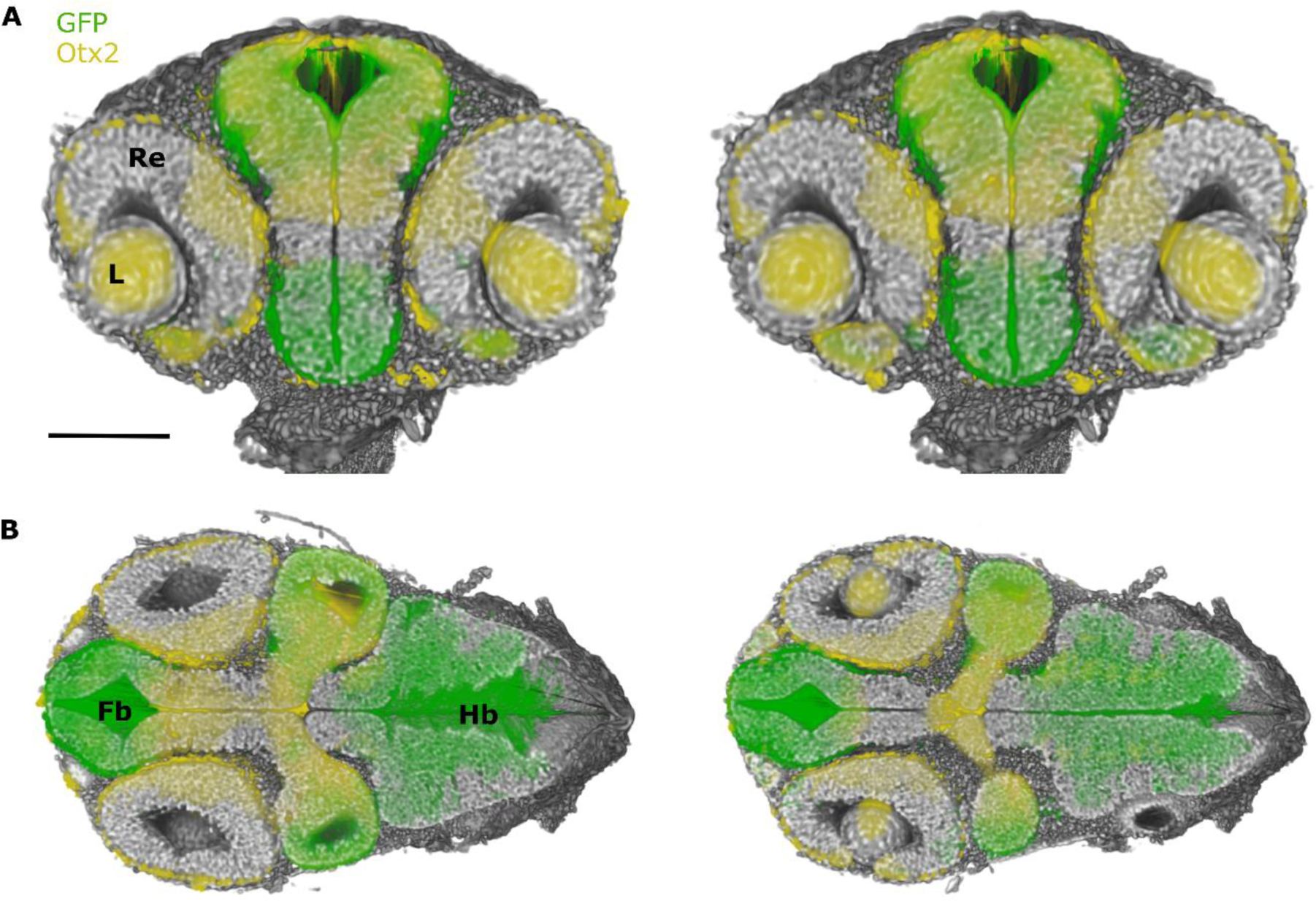
Characterisation of *Sox2(600bp)::GFP line* at stage 28 by multimodal approach. (A) Retinal domains outlined by Otx2 expression only and non-retinal neuronal domains expressing *sox2* and Otx2 in axial cuts along the eye and eye with optical nerve. (B) Specificity of *sox2* expression to the neuronal non-retinal domain in coronal sections of the brain region. Organs are labelled as follows: lenses (L), retina (Re), forebrain (Fb), and hindbrain (Hb). The scale bar is 100 µm.

## Video legends

**Video 1 Neuronal genes in the context of tissue anatomy of the medaka embryo stage 24 visualised by the multimodal approach.** 3D rendering of all contrast modalities, see Figure 2, Otx2 in yellow, PH3 in cyan and HuC/D in magenda visualised with virtual cuts through the whole volume.

**Video 2 Tissue architecture of phenotypic variations in *Rx3^saGFP^* mutants based on X-ray tomography.** 3D renderings of segmented tissues, that is brain (blue), retina (green), and lens (orange) in wild-type, incomplete and complete phenotypes.

**Video 3 Complex 3D network of the lumen in medaka-derived organoids revealed by X-ray tomography.** 3D rendering of organoid and segmented lumen presented as surface view in green within it, see also Figure 5D.

